# Seasonal variability in copepod biomass in a cyclonic eddy in the Bay of La Paz, southern Gulf of California, Mexico

**DOI:** 10.1101/2020.03.13.990382

**Authors:** Franco Antonio Rocha Díaz, María Adela Monreal Gómez, Erik Coria Monter, David Alberto Salas de León, Elizabeth Durán Campos

## Abstract

As one of the main groups composing marine zooplankton, copepods play an important role due to the position they occupy in the trophic web. Study of their biomass and relationship with the physical conditions of the water column are essential in order to evaluate the trophic structure and functions of any aquatic ecosystem. As a contribution to this topic, we assessed the copepod biomass inside a cyclonic eddy system during two different seasons in the Bay of La Paz in the southern Gulf of California, a region characterized by high biological productivity. Two oceanographic expeditions took place in the winter of 2006 and summer of 2009 on which a conductivity-temperature-depth (CTD) probe was used to determine the physical structure of the water column and oblique zooplankton hauls collected zooplankton samples. Satellite data were used to visualize chlorophyll-a distribution patterns. The results showed the presence of a well-defined mesoscale cyclonic eddy in both seasons, with high chlorophyll-a (CHLA) values at the edges of the eddy. Maximum values for copepod biomass were observed in winter and their distribution corresponded well with the circulation pattern and the CHLA values, forming a belt shape following the periphery of the eddy. The results presented herein highlight the impact of the mesoscale eddy on the planktonic ecosystem through its influence on hydrographic conditions in the water column. Other factors, such as ecological interactions, population dynamics, and feeding habits may play a role as well. Feeding behavior in particular is affected by high CHLA concentrations observed around the eddy which represent a source of food for these organisms.

## Introduction

Copepods are the most abundant multicellular organisms on Earth [1], [2] and in the marine ecosystem where they are of prime importance due to the position they occupy in the trophic web [3]. As mostly herbivorous organisms, they feed on phytoplankton and therefore represent a link between lower and higher trophic levels. Several species of copepods have high commercial value [4]. Additionally, trophic web dynamics contribute to the removal of CO2 from the atmosphere, through sedimentation of inorganic and organic carbon compounds included in fecal pellets, and to the appropriate functioning of the biological or carbon pump [3].

Typically, zooplankton biomass is indicative of secondary production and estimation of this parameter is essential to evaluate trophic structure and function in any aquatic ecosystem [5]. Changes in zooplankton biomass are closely related to several factors, including variations in the salinity field [6], the temperature regime [7], and the availability of food [8].

Another source of variability in zooplankton biomass is the presence of hydrodynamic processes, which modify the hydrographic structure of the water column and exert a remarkable effect on productivity by introducing nutrients into the euphotic zone, with consequent enhancement in phytoplankton biomass due to the increased availability of food for the zooplankton community [9]. These hydrodynamic processes are present throughout the water column at different scales, including internal waves, fronts, and eddies [10].

Mesoscale eddies (radii 10–100 km) are high energy hydrological structures of prime importance in any marine ecosystem [11]. These structures, are recognized as cyclonic, anticyclonic, and mode-water with a noticeable impact on the planktonic ecosystem [12]. It has been demonstrated that the presence of cyclonic eddies modulate the structure and biomass of the zooplankton community in different parts of the world, including the Mediterranean Sea [13], the Sargasso Sea [14], the Madagascar Channel [15], the Pacific Ocean off central-southern Chile [16], and the Hudson Bay in Canada [9].

Although the impact of mesoscale eddies on zooplankton biomass has been relatively well demonstrated worldwide, uncertainties remain about the role of these structures on particular groups of zooplankton, in this case copepods. As a contribution to this topic, this research aimed to assess the biomass of copepods in a mesoscale cyclonic eddy system in the Bay of La Paz, Gulf of California. Hydrographic data and zooplankton samples were collected during two oceanographic expeditions in two contrasting seasons, summer 2009 and winter 2006. We hypothesized that the copepod biomass will vary with respect to season and hydrodynamics of the mesoscale eddy system. It is our intention that this study contributes to a better understanding of the influence of mesoscale eddies on particular and pivotal zooplankton groups, such as copepods, in one of the most productive marine ecosystems in the world.

## Materials and methods

The Bay of La Paz is the largest basin in the Gulf of California. The region is highly dynamic, with a wide seasonal and interannual variability due to atmosphere-ocean interactions. In winter, the prevailing winds are predominantly northwesterly, with high and persistent speeds exceeding 10 m s^-1^. In summer, southeasterly winds blow at approximately 5 m s^-1^, with frequent calms [17].

The study area is recognized for its high biodiversity which has been linked to the hydrodynamics of the region, involving the presence of a quasi-permanent cyclonic eddy which exerts a great impact on the planktonic ecosystem by supporting production at high trophic levels. Indeed, this cyclonic eddy promotes a nutrient-rich Ekman pump that fertilizes the euphotic layer [18] and induces a differential distribution in phytoplankton between diatoms and dinoflagellates [19] and a differential aggregation of zooplankton inside the eddy field [20], which then impacts the whole pelagic food web.

In this research, high-resolution hydrographic data and zooplankton samples were obtained during two research cruises onboard the R/V “El Puma” (UNAM) in winter (February 22–27, 2006) and summer (August 11–20, 2009). Using a CTD probe (SeaBird 19 plus) previously calibrated by the manufacturer, we acquired data at 45 hydrographic stations across the bay (Fig. 1b). The CTD probe was lowered at a rate of 1 m s^-1^ and configured to store data at 24 Hz. Immediately following the CTD cast, oblique zooplankton hauls were performed using a bongo system with a 60 cm diameter mouth equipped with a net mesh size of 333 µm. The haul time was 15 min at 1 m s^-1^ at 13 stations in winter and 9 stations in summer (Fig. 1b); in both cases, the stations were located in Alfonso Basin and in the northern reaches of the bay near its connection to the Gulf of California. The selection of stations in Alfonso Basin was made in consideration of previous researches that revealed the presence of a quasi-permanent mesoscale cyclonic eddy [17]. Zooplankton organisms were collected between depths of 200 m and the surface. The water volume filtered during the haul was calculated using calibrated flowmeters (General Oceanics Inc.) placed in each net. Once on board, the samples were fixed for 24 h with a solution of 4% formalin plus sodium borate, then preserved in 70% ethanol.

**Fig 1.**
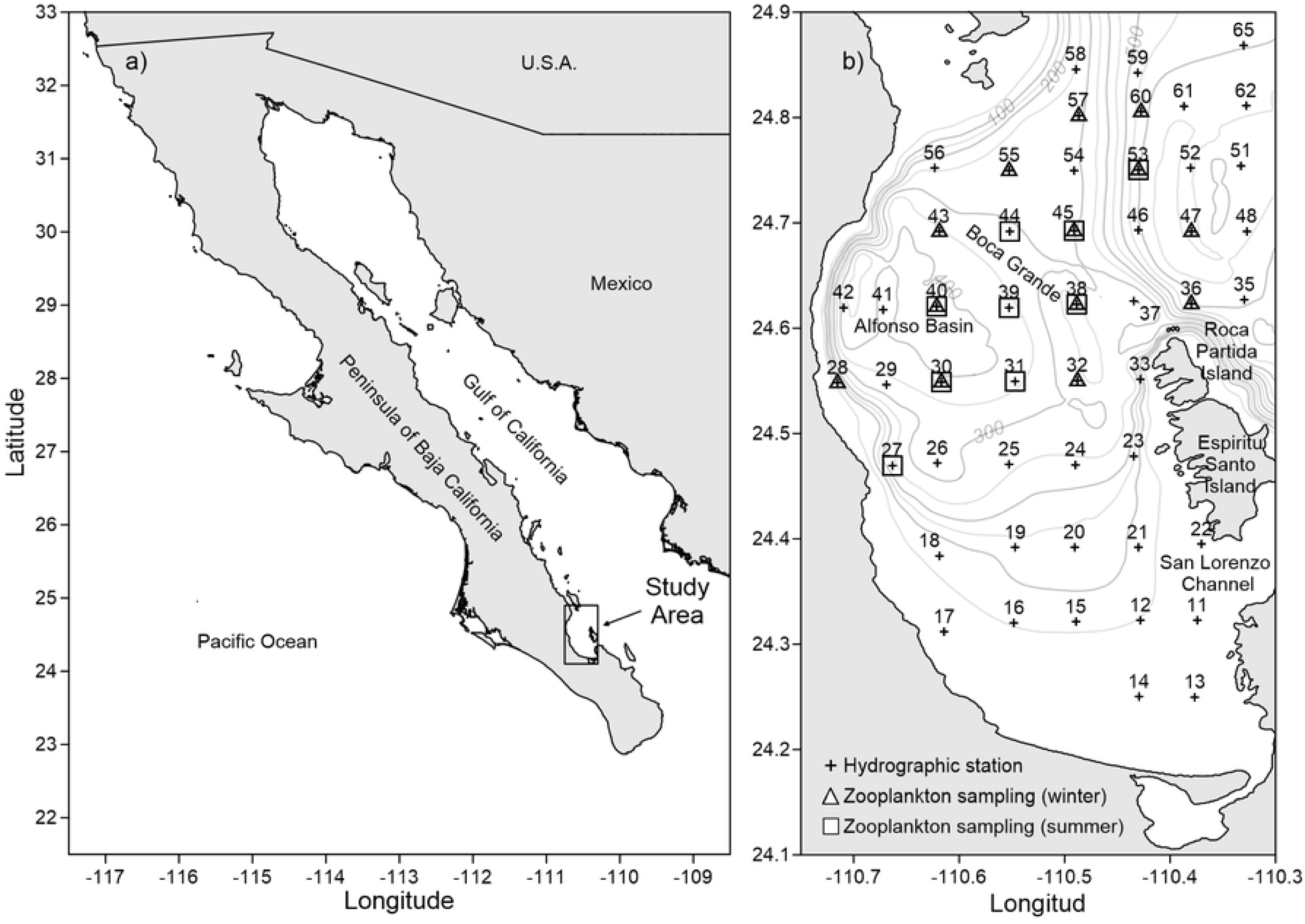
Location of the Gulf of California and Bay of La Paz. Location of: a) Gulf of California and b) Bay of La Paz. Bathymetry (m), CTD cast (+), zooplankton samples in winter (Δ), zooplankton samples in summer (□).

The CTD data were processed following the subroutines provided by the manufacturer (SBE Data Processing V.7.26.7 software) averaged over 1 dbar intervals. Then the conservative temperature Θ (°C), absolute salinity SA (g kg^-1^), and density *σ* (kg m^-3^) were derived following the algorithms of the Thermodynamic Equation of Seawater-2010 (TEOS-10) [21]. Geostrophic circulation patterns during the two research cruises were obtained by means of the geostrophic method for calculating velocity relative to the bottom [22]. These circulation patterns were analyzed at the base of the thermocline, which was obtained by the maximum vertical gradient method 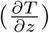.

Satellite images of the CHLA concentration, as an indicator of phytoplankton biomass, with a resolution of 1 km/pixel were obtained from the NASA Ocean Biology Processing Group (OBPG) concurrent with the dates of both research cruises. The images were processed using levels 1 and 2 with SeaDAS software provided by OBPG. Different masks/flags (STRAYLIGHT, CLDICE, LAND, and HILT) were applied in order to purge bad or low-quality data when the images were generated.

In the laboratory, the zooplankton samples were sequentially split with a Folsom mechanism and individual copepods were identified following Boltovskoy [23], counted, and then separated using a Carl Zeiss stereo dissecting microscope. The copepods picked from the samples were pooled in a glass Petri dish and separated into three groups: calanoid copepods, cyclopoid copepods, and all copepodite stages. In order to quantify the biomass for each group (wet weight, mg/100 m^3^) at each station, the ethanol was removed using a Millipore system manually pumped through pre-weighed nitrocellulose membrane filters (0.45 *µ*m, 47 mm diameter, Millipore Corp., USA). Then, using an analytical balance (Sartorius BP211D, resolution: 0.1 mg/210 g), differences in weight were obtained. Finally, biomass (mg/100 m^3^) was obtained following the protocols described in Duran-Campos et al. [20], [8].

## Results and Discussion

The geostrophic circulation patterns were analyzed at the base of the thermocline. In the winter (2006) the mixing layer was 50 m deep; in summer (2009), it was 30 m deep. In both cases, current patterns were characterized by the presence of a mesoscale cyclonic eddy located in Alfonso Basin; however, the intensity and shape of the eddy showed differences between seasons. During the winter, the results showed the mesoscale eddy with a diameter of 30 km, reaching a velocity of 50 cm s-1 at its periphery and occupying the deepest part of the bay and the Boca Grande region at its confluence with the Gulf of California, where exchanges between basins occur. In this region, the velocities showed an incremental increase and the circulation pattern bifurcates into the bay and to the north (Figure 2a–c). The geostrophic velocities at 30 m depth (not shown) in the mixing layer were similar to those at 50 m depth. The circulation also showed a clockwise current in the southwestern region of the bay close to the coast. During summer (2009) at 30 m depth, the geostrophic velocities showed the presence of a well-defined mesoscale cyclonic eddy occupying the same area as in winter (2006) of similar diameter; however, at the base of the thermocline (30 m depth), the geostrophic velocities reached values of 70 cm s-1 at the periphery of the eddy (Figure 2d–f).

**Fig 2.**
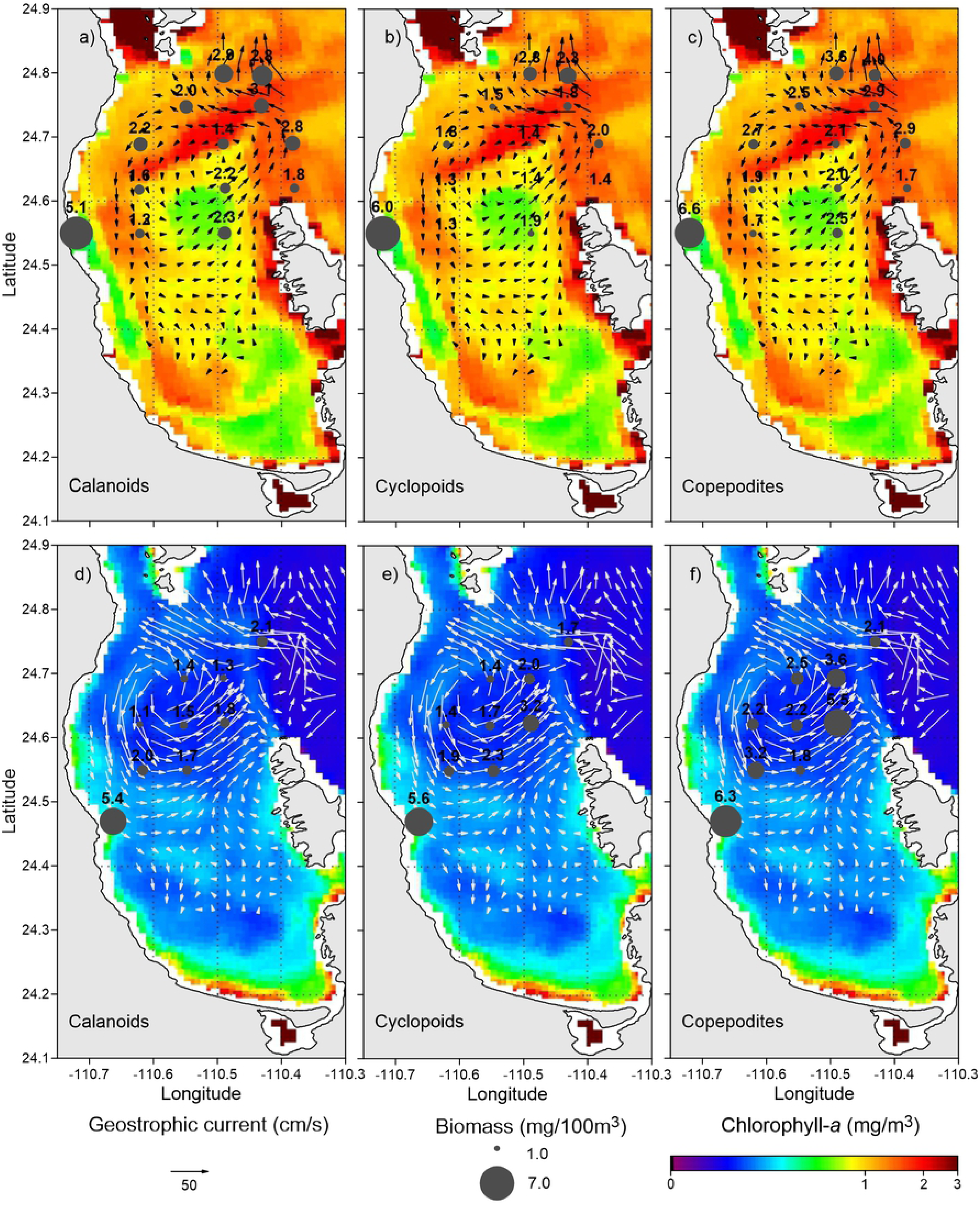
Geostrophic current and surface chlorophyll-a distribution. Top panel: Geostrophic current pattern at the base of the thermocline (50 m), surface chlorophyll-a distribution (mg m^-3^), and biomass (mg/100 m^3^) of: a) calanoid copepods, b) cyclopoid copepods and c) all copepodite stages (winter 2006). Bottom panel: Geostrophic current pattern at the base of the thermocline (30 m), surface chlorophyll-a distribution, and biomass (mg/100 m^3^) of: d) calanoids, e) cyclopoids and f) all copepodite stages (summer 2009).

The results of CHLA concentration (as an indicator of phytoplankton biomass) obtained by satellite showed a remarkable coupling with the circulation pattern. During the winter of 2006, an interesting pattern of distribution was observed in the northern bay in connection with the Gulf of California where high values (3 mg m^-3^) formed an enhanced area with a belt shape around the cyclonic eddy that gradually decreased towards the center (Fig. 2 a–c); during the summer of 2009, the CHLA concentration was lower than in winter, showing highest values in the southern coastal region with a secondary peak (0.8 mg m^-3^) forming a circular shape around the cyclonic eddy (Figure 2 d–f).

In order to establish the coupling between the circulation patterns obtained at the time of our observations of CHLA concentrations and the copepods studied (calanoids, cyclopoids, and all copepodite stages), the results of the biomass for each group were superimposed (Fig. 2). The results showed a pattern of distribution with progressive changes from the connection with the Gulf of California to the interior of the bay, as well as from the periphery to the center of the cyclonic eddy.

In winter, biomass concentrations were slightly higher than those observed in summer, showing two regions of maximum biomass for the three target groups: one located at the junction with the gulf, one in the Boca Grande region, and a third in the western region close to the coast (*>* 5 m/100 m^3^) (Fig. 2 a–c). A pattern of change was observed within the eddy field with high values associated with the periphery of the eddy and defining its circumference. During the summer, the highest values, located in the western region close to the coast (*>* 5 mg/100 m^3^), coincided with secondary high values around the periphery of the eddy (Fig. 2 d–f).

These results showed that the copepod groups were influenced by the cyclonic eddy through their actions on the hydrographic conditions, and possibly as a result of several additional processes such as ecological interactions, population dynamics, and feeding habits, particularly by the high CHLA concentrations observed around the eddy which potentially represent a source of food for these organisms. This could also be related with bottom up mechanisms, where high trophic levels (e.g., fishes) could use this region to feed. High CHLA concentrations around the eddy are attributed to mechanisms of fertilization by nutrients documented in these structures along with the mixing processes that occur on the periphery, ensuring the availability of nutrients for the phytoplankton communities, and thus for the zooplankton grazers [24]. In particular, copepods are one of the most successful organisms in the marine environment because their torpedo shape enhances mobility which is propitious for locating food [2]. Along this line of thought, the high copepod biomass values found in the three groups analyzed near the connection with the Gulf of California can be explained by the presence of a bathymetric sill where important processes take place (e.g., hydraulic jumps) that fertilize the euphotic zone [25]. The highest biomass was observed in the western portion of the bay close to the coast, and can be explained by the presence of a phosphate mining industry that fertilizes the region, causing phytoplankton blooms that increase the herbivorous zooplankton population. The progressive changes observed in copepod biomass around the eddies could be induced by the advection generated by convergent movements induced by the cyclonic structure.

The low concentration of CHLA observed in the center of the eddy in both seasons could be related to the predominance of certain heterotrophic phytoplankton groups (e.g., dinoflagellates), while the high copepod abundance associated with high CHLA values in the form of a belt-shaped area are related to the predominance of diatoms, as previously reported by [19].

Changes observed in phytoplankton biomass according to season could be associated with normal periods of heating and cooling of surface layers in summer and winter, which induce mixing in winter, thus increasing the concentration of nutrients leading to high CHLA values [26] available for the zooplankton. Martínez-López et al. [27] also reported this pattern of high biological productivity during winter in Alfonso Basin.

## Acknowledgments

Ship time for both research cruises onboard the R/V ‘El Puma’ was funded by Universidad Nacional Autónoma de Mexico. CONACYT, Mexico sponsored FARD through a graduate scholarship. We appreciate the assistance of the captain and crew for assistance in seagoing activities, as well as the scientific staff on both cruises. We thank the NASA Ocean Biology Processing Group for the satellite products used in this study. Sergio Castillo Sandoval provided technical support during the laboratory analyses and Jorge Castro improved the figures.

